# Extreme genome selection towards complete antimicrobial resistance in a nosocomial strain of *Stenotrophomonas maltophilia* complex

**DOI:** 10.1101/735555

**Authors:** Sanjeet Kumar, Kanika Bansal, Prashant P. Patil, Amandeep Kaur, Satinder Kaur, Vikas Gautam, Prabhu B. Patil

## Abstract

We report first complete genome sequence and analysis of an extreme drug resistance (XDR) nosocomial *Stenotrophomonas maltophilia* that is resistant to the mainstream drugs i.e. trimethoprim/sulfamethoxazole (TMP/SXT) and levofloxacin. Taxonogenomic analysis revealed it to be a novel genomospecies of the *Stenotrophomonas maltophilia* complex (Smc). Comprehensive genomic investigation revealed fourteen dynamic regions (DRs) exclusive to SM866, consisting of diverse antibiotic resistance genes, efflux pumps, heavy metal resistance, various transcriptional regulators etc. Further, resistome analysis of Smc clearly depicted SM866 to be an enriched strain, having diversified resistome consisting of *sul1* and *sul2* genes. Interestingly, SM866 does not have any plasmid but it harbors two diverse super-integrons of chromosomal origin. Apart from genes for sulfonamide resistance (*sul1* and *sul2*), both of these integrons harbor an array of antibiotic resistance genes linked to ISCR (IS91-like elements common regions) elements. These integrons also harbor genes encoding resistance to commonly used disinfectants like quaternary ammonium compounds and heavy metals like mercury. Hence, isolation of a novel strain belonging to a novel sequence type (ST) and genomospecies with diverse array of resistance from a tertiary care unit of India indicates extent and nature of selection pressure driving XDRs in hospital settings. There is an urgent need to employ complete genome based investigation using emerging technologies for tracking emergence of XDR at the global level and designing strategies of sanitization and antibiotic regime.

**Impact Statement:** The hospital settings in India have one of the highest usage of antimicrobials and heavy patient load. Our finding of a novel clinical isolate of *S. maltophilia* complex with two super-integrons harbouring array of antibiotic resistance genes along with antimicrobials resistance genes indicates the extent and the nature of selection pressures in action. Further, the presence of ISCR type of transposable elements on both integrons not only indicates its propensity to transfer resistome but also their chromosomal origin suggests possibilities for further genomic/phenotypic complexities. Such complex cassettes and strain are potential threat to global health care. Hence, there is an urgent need to employ cost-effective long read technologies to keep vigilance on novel and extreme antimicrobial resistance pathogens in populous countries. There is also need for surveillance for usage of antimicrobials for hygiene and linked/rapid co-evolution of extreme drug resistance in nosocomial pathogens. Our finding of the chromosomal encoding XDR will shed a light on the need of hour to understand the evolution of an opportunistic nosocomial pathogen belonging to *S. maltophilia*.

**Repositories:** Complete genome sequence of *Stenotrophomonas maltophilia* SM866: CP031058

## INTRODUCTION

According to World health organization (WHO) list *Stenotrophomonas maltophilia* is one of the leading opportunistic multidrug-resistance pathogen in hospitals worldwide (http://www.who.int/drugresistance/AMR_Importance/en/). *S. maltophilia* causes variety of infections including respiratory tract infections, bloodstream infections, bone and joint infections, urinary tract infections, endocarditis and meningitis (1). *S. maltophilia* is having intrinsic resistance to multiple groups of widely used antibiotics such as: cephalosporins, carbapenems, aminoglycosides and macrolides (2). Till now, trimethoprim/sulfamethoxazole (TMP/SXT) and levofloxacin were considered as the mainstream therapy for such infections (3, 4). Elevated reports of resistance to both of these drugs is a growing concern worldwide (1). Hence, their rapid identification and investigation are key towards the successful treatment.

*S. maltophilia* has been reported to acquire resistance through integrons, transposons and plasmids conferring multiple resistance genes (5). There are various PCR-based or draft genome based studies for the identification of these resistant strains (6–8). Until now, ISCRs (IS91-like elements common regions), considered as gene-capturing systems of 21^st^ century are also found in the range of extreme drug resistant (XDR) bacteria carrying antibiotic and antimicrobial resistance genes (9, 10). For instance, in *Shewanella xiamenensis* the ISCR was linked with antibiotic resistance genes along with resistance genes for biocides heavily used in hospital settings such as: quaternary ammonium compounds (*qac*), heavy metals (mercury resistance genes) etc. (11). Interestingly, ISCR in *S. maltophilia* strains are also associated with *sul* genes. These are majorly reported from plasmids or with very low confidence, predicted to be chromosomally encoded (5, 12). Lack of complete genome based studies of XDR *S. maltophilia*, limits our understanding of the genomic context of the resistance genes cassettes. Such an information will be critical to understand genome dynamics particularly for new strains carrying multiple alleles that are spreading by integrons which carry array of antibiotic resistance genes linked to ISCR elements (13).

Developing countries like India have high selection pressure because of heavy antibiotic and biocide usage and high patient density coupled with tropical climate setting. Recent reports revealed rapid emergence of extreme drug resistance in India in sulfonamide resistant *S. maltophilia* and carbapenem resistance conferred by New Delhi metallo-beta-lactamase 1 (NDM-1) in *Enterobacteriaceae* strains (1, 14). Furthermore, studies have reported emergence of biocides (absolute ethanol, povidone iodine, sodium hypochlorite, and QACs) resistance among multidrug resistance bacteria such as: *Escherichia coli, Klebsiella pneumoniae, Pseudomonas aeruginosa* and *Staphylococcus aureus* and diverse biofilm forming bacteria of dairy niche (15, 16). Use of emerging cost-effective nanopore technologies are revolutionizing investigation of such extreme antimicrobial resistance by allowing rapid access of complete genome information. In the present study, we report isolation and complete genome based investigation of a novel sequence type (ST-366) of extreme drug resistant strain belonging to Smc from a tertiary care center in northern India. Complete genome level study of the strain indicates its highly dynamic nature and found to harbor fourteen dynamic regions (DRs). The strain also harbours two diverse chromosomal integrons linked to an array of antibiotic resistance genes along with IS elements and antimicrobial resistance genes. Overall, the study indicated serious long term consequences of high selection pressure for rapid emergence and evolution towards complete antimicrobial resistance strains in Indian hospitals. At the same time, the study highlights the need to employ rapid and long read technologies to carry out complete genome based investigations to understand and manage extreme drug resistance at global level.

## METHODS

### Bacterial isolation, identification and antimicrobial susceptibility testing

Bacterium was isolated from the respiratory specimen of hospitalized patient from Postgraduate Institute of Medical Education and Research (PGIMER), Chandigarh in 2013. Ethics approval and each patient’s written consent was not required as it was a part of routine clinical testing. Bacterial cultures were grown in the nutrient broth at constant shaking of 200 rpm at 37 °C. DNA isolation was performed with ZR fungal/ bacterial DNA MiniPrep Kit (Zymo research) in accordance to manufacturer instructions. In order to identify the bacterial species partial 16S rRNA sequencing was performed using genomic DNA. Primers used for 16S rRNA amplification were 27F and 1492R. For Bacterial identification, EZ Bioclould (17) (http://www.ezbiocloud.net/identify) was used. Further, antimicrobial susceptibility testing was performed by the Kirby-Bauer disc diffusion method as per the Clinical Laboratory Standards Institute (CLSI), 2017 guidelines against trimethoprim-sulfamethoxazole (25 mcg), chloramphenicol (30 mcg), levofloxacin (5 mcg) and ceftazidime (30 mcg) antibiotics.

### Genome sequencing, assembly and annotation

a. **Nanopore sequencing**: Bacterial cultures were grown overnight in 15 ml of Luria-Bertani broth. Bacterial cells were harvested and high quality bacterial genomic DNA was isolated using Qiagen DNeasy Blood & Tissue Kit. DNA was purified using 0.45x Agencourt AMPure XP beads (Beckman Coulter, High Wycombe, UK) and quantified using Nanodrop 1000 v3.8 (Thermo Fisher Scientific) and Qubit 2.0 (Thermo Fisher Scientific). Purified DNA sample was end-repaired (NEBnext ultra II end repair kit, New England Biolabs, MA, USA), cleaned up with 1x AMPure XP beads (Beckmann Coulter, USA). Native barcode ligation was performed with NEB blunt/ TA ligase (New England Biolabs, MA, USA) and cleaned with 0.5x AMPure XP beads. Qubit quantified and barcode ligated DNA samples were pooled at equimolar concentration to attain 1 µg pooled sample. Adapter ligation (BAM adapters) was performed for 15 minutes using NEBNext Quick Ligation Module (New England Biolabs, MA, USA). Library mix was cleaned up using 0.4X AMPure XP beads (Beckmann Coulter, USA) and finally sequencing library was eluted in 15 µl of elution buffer.
b. **Illumina Sequencing**: Genomic DNA was extracted from SM866 using ZR Fungal/Bacterial DNA MiniPrep Kit (Zymo Research, Irvine, CA, USA) and quantified using Qubit 2.0 Fluorometer (Life technologies). Short reads libraries was sequenced on Illumina Miseq (Illumina, Inc., San Diego, CA, USA). Illumina sequencing library was prepared using Nextera XT sample preparation kit (Illumina, Inc., San Diego, CA, USA) with dual indexing adapters and sequencing using 250*2 paired end sequencing kit. The adapters were trimmed while sequencing by internal software Illumina MiSeq Control v.2.5.0.5 (Illumina, Inc., San Diego, CA, USA).

Genome assembly was first performed using the Unicycler v0.4.6 (18) with ONT long reads with bold mode. The complete genome obtained using ONT reads was then polished for multiple rounds of Pilon v1.22 (19) using Illumina raw reads of the sample SM866. The genome was assembled in a closed circular single chromosome. Annotation was performed using NCBI-PGAP (20) annotation pipeline. Assembly quality in terms of completeness and contamination was accessed using the CheckM v1.0.11 (21). Genome coverage of the SM866 was calculated using BBMap v36.92 (22).

#### 16S rRNA gene Phylogeny

Complete 16S rRNA gene sequence from the type strains of the genus *Stenotrophomonas* was fetched from their corresponding INSDC number (http://www.insdc.org/) from LPSN (23) given for their species definition (http://www.bacterio.net/stenotrophomonas.html). 16s rRNA sequence from SM866 was fetched from complete assembled genome using standalone version of RNAmmer v1.2 (24). All the complete 16S rRNA gene sequences were aligned using ClustalW (25). Phylogenetic tree was constructed based on neighbor-joining method was created with 1000 bootstrap. *Xanthomonas campestris* ATCC33913 (26) was used as outgroup to genus *Stenotrophomonas*.

#### Multi-locus sequencing typing (MLST)

Sequence type analysis of the strain SM866 was done using center for genomic epidemiology MLST 2.0 (https://cge.cbs.dtu.dk/services/MLST/) and PubMLST (https://pubmlst.org/bigsdb).

### Phylogenomic and taxonogenomics based identification and characterization

More robust phylogeny was constructed using > 400 conserved gene using PhyloPhlAn (27). Complete genome proteome of all the type species of genus *Stenotrophomonas* was used for the construction of the phylogenetic tree. *X. campesteris* ATCC 33913 was used as outgroup. Further, taxonogenomic analysis of the strain was performed using orthoANI (28) and dDDH values (29).

### Dynamic regions and resistome analysis

The unique DRs of SM866 were detected using BRIG v0.95 (30) which were further confirmed using blastn package of BLAST++ (31). A well characterized nucleotide sequence of the antibiotic resistance gene clusters and efflux pumps were retrieved from the complete genome of *S. maltophilia* K279a (32). A standalone blast was performed with these gene as query with SM866 and the previously reported strains of Smc (33). A heat map showing the presence and absence of the gene cluster was generated using GENE-E v3.0.215 (http://www.broadinstitute.org/cancer/software/GENE-E). Further, schematic diagram of the cassettes I and II were generated using Easyfig v2.2.2 (34).

### Plasmid profiling

The plasmid profiling was performed for isolate SM866 using FosmidMAX DNA Purification kit (Epicenter, Illumina). For this the bacterial sample was grown in 10 ml of LB and incubated for 28 °C for 16 hours. Bacterial cells were harvested by centrifugation at 8000 rpm for 5 minutes. Plasmid isolation was performed as per the manufacturer recommendations. 5µl of sample was loaded on 0.7% agarose gel to check the presence of plasmid (Supplementary figure 2). To reconfirm the absence of plasmid, Plasmid-Safe(tm) ATP-Dependent DNase was performed in accordance to manufacturer recommendation and *S. maltophilia* ATCC13637 (T) was used as a control.

## RESULTS

### Bacterial isolation, identification and antimicrobial susceptibility

The strain SM866 was isolated in 2013 from respiratory specimen from an intensive care unit patient admitted in a tertiary care hospital in Northern India (20). SM866 exhibited 99.5% partial 16S rRNA nucleotide sequence identity with the type strain *S. maltophilia* ATCC 13637(T). The 16S rRNA gene phylogeny using all the type strains of the genus *Stenotrophomonas* clearly showed that our isolate was in Smc (supplementary figure 1(A)). For the phylogeny, we have considered all the known novel genomospecies of the Smc (SM5815, SM3123, SM10507 and SM16975) (33). Further, antibiotic resistance profiling revealed that SM866 is resistant to all the antibiotics tested which includes trimethoprim/ sulfamethoxazole, levofloxacin, chloramphenicol and ceftazidime. The zone of inhibition for the antibiotics used is in table 1.

**Table 1:** SM866 zone of inhibitions for antibiotics: TMP-SMX: trimethoprim-sulfamethoxazole (25mcg), CHL: chloramphenicol (30 mcg), LEV: levofloxacin (5 mcg) and CAZ: ceftazidime (30 mcg).

| Antibiotics | SM866 (zone diameter in mm) | Result interpretation |  |  |
| --- | --- | --- | --- | --- |
| | | (≥) Sensitive (S) (in mm) | Intermediate (I) (in mm) | (≤) Resistant (R) (in mm) |
| TMP-SMX | 0 (R) | 16 | 11 to 15 | 10 |
| CHL | 0 (R) | 18 | 13 to 17 | 12 |
| LEV | 11 (R) | 17 | 14 to 16 | 13 |
| CAZ | 0 (R ) | 21 | 18 to 20 | 17 |

### Multi-locus sequence typing and population structure analysis

Multilocus sequence typing of SM866 revealed that out of seven housekeeping genes (*atpD, gapA, guaA, mutM, nuoD, ppsA* and *recA*), it harbors three novel housekeeping alleles i.e. *guaA, mutM* and *nuoD* with newly assigned accession numbers: 272, 147 and 141 respectively. Hence, SM866 was assigned a novel sequence type, ST-366. Further, population structure analysis was carried out considering all the STs of *S. maltophilia*, clearly depicting that SM866 is a singleton ST and it does not belong to clonal complex of any of the already known STs (supplementary figure 1(B)).

### Genome sequencing and annotation

The complete circular genome of SM866 with one chromosome (and no plasmid) of 5.08 MB with 60X coverage and 66.03% GC content was obtained by assembly of the Oxford Nanopore long reads and polishing with Illumina short reads. SM866 genome has been submitted to the public repository NCBI with accession number CP031058. PGAP annotation resulted in 4,715 CDS, 73 tRNA and 4 ncRNAs. Further, CheckM estimated 99.76% completeness and 0.72% contamination in the assembled genome (https://figshare.com/s/72aee1707e6a6182ac6d). Further, plasmid profile of the strain SM866 was confirmed by plasmid screening resulted in no plasmid detection in SM866 and the *S. maltophilia* ATCC13637 (T) (supplementary figure 2(A)). Although, there was presence of very faint band in SM866, which when treated with ATP-dependent DNase method was degraded, inferring that to be a genomic DNA contamination (supplementary figure 2(B)). Hence, this confirms that SM866 is devoid of plasmid, as also indicated by complete genome.

### Phylogenomic and taxonogenomic analysis

Taxonogenomic analysis of SM866 was carried out to clearly know its species status. ANI and dDDH values for SM866 were below the cut-off values for species delineation. Within Smc, values ranged from 87%-93% and 34%-48% for ANI and dDDH respectively (figure1). Hence, this analysis suggested SM866 to be a novel genomospecies in Smc. Furthermore, phylogenomic analysis done by taking *Xanthomonas campestris* ATCC33913 (T) as an outgroup suggested that SM866 falls within the Smc complex (figure 1).

**Figure 1:**
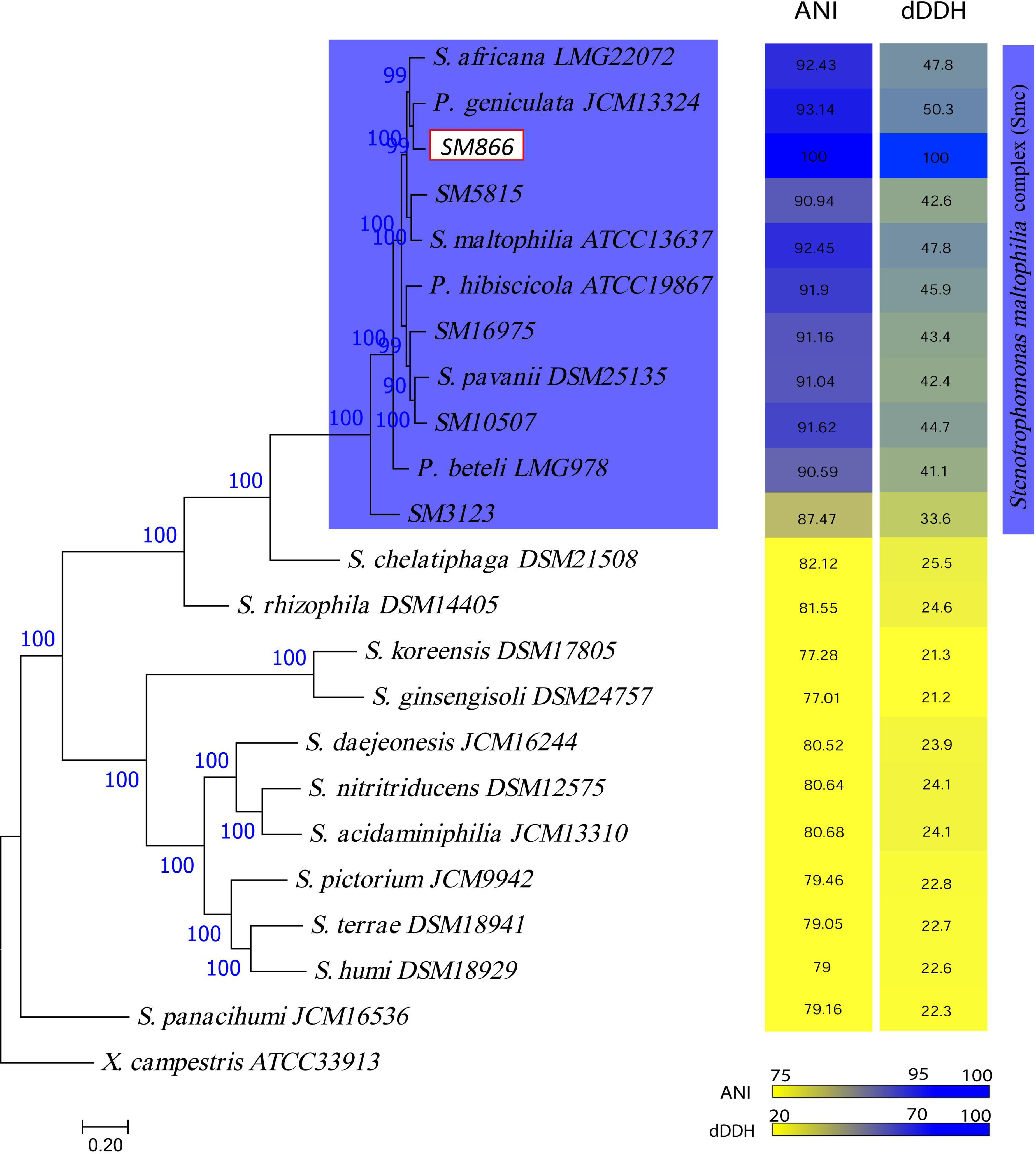
**PhyloPhlAn phylogeny and taxonogenomic analysis** of representative strains of *Stenotrophomonas* genus along with SM866 (in red box) and Smc is highlighted in blue box. ANI and dDDH values for all the representative strains taking SM866 as reference are shown in the heat maps next to the tree generated on the scale as shown in the figure.

### Understanding genome dynamics in evolution of extreme drug resistance

SM866 was found to be trimethoprim/sulfamethoxazole and levofloxacin resistant, unlikely of previous genomospecies of the Smc. We focused on the unique regions to SM866, which were absent from other Smc complex strains. Interestingly, we found 14 dynamic regions (DRs) exclusively in SM866 (supplementary table 1). Functional classification of the DRs along with their genomic locations is provided in the figure 2 and supplementary table 1. Here, a range of antibiotic resistance genes (tetracycline, chloramphenicol, beta-lactamases, sulfonamide-resistance, aminoglycosides), efflux pumps (RND, ABC, cation transporter, Na+/H+ antiporter), heavy metal resistance (copper, mercury, arsenate), quaternary ammonium compound (QacE), glycosyltransferases, methyltransferases, acetyltransferases, DEAD/DEAH box helicase, disulfide bond formation protein DsbA, DNA repair protein, transcriptional regulators (MerR, ArsR, XRE, CopG, Lrp/AsnC, LysR, TolC, TetR/AcrR, LuxR), toxin-antitoxin system HipA, lipoproteins, lysozyme, redox-sensitive transcriptional activator SoxR, integrases, sensor kinases, response regulators, IS elements, phage proteins, transposases, conjugal transfer proteins (TrbBCDEFGIJLK, TraGJ) and proteases etc. were found in the DRs unique to SM866.

**Figure 2:**
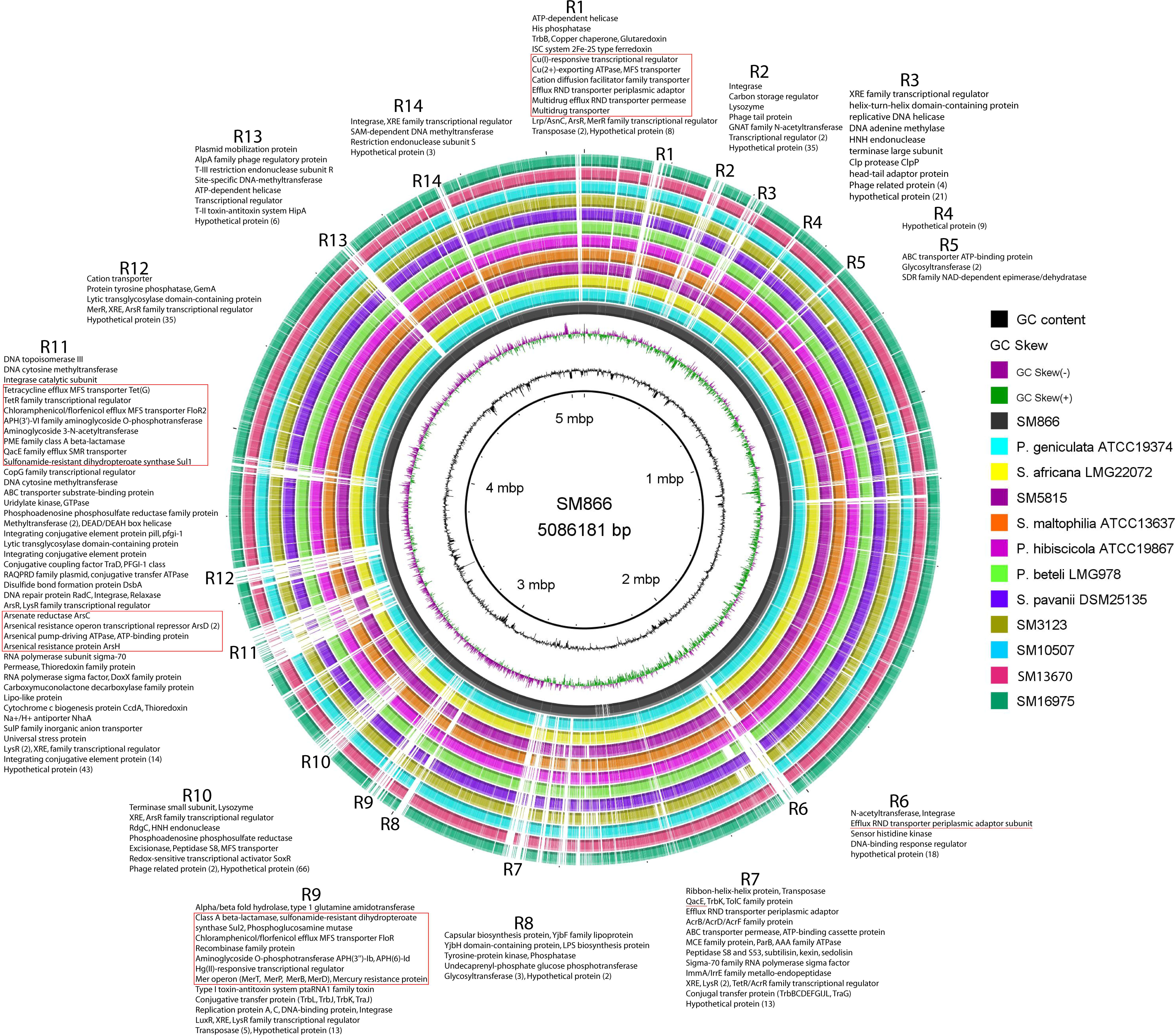
**Circular representation of SM866 along with other Smc genomospecies.** Rings are color coded for different strains. The innermost two rings represent GC content (%) and GC skew. Here, R1-R14 are DRs exclusively present in SM866, annotation for the regions are indicated adjacent to regions and genomic locations of the DRs is provided in supplementary table 1.

### Resistome analysis of Smc genomospecies

Resistome of Smc genomospecies is represented in figure 3. Here, among all the Smc genomospecies, only SM866 was found to have *sul1* and *sul2* genes. Here, drug resistance genes for aminoglycosides, beta-lactams, chloramphenicol, fluoroquinolones; efflux pumps like RND, SMR, MFS, MATE and ABC transporters were examined. This reveals the underlying genes for intrinsic antibiotic resistance of the Smc strains. Interestingly, in the vast resistome of SM866, *sul1* and *sul2* are exclusively present in SM866 only, which have been also detected in R11 and R9 respectively.

**Figure 3:**
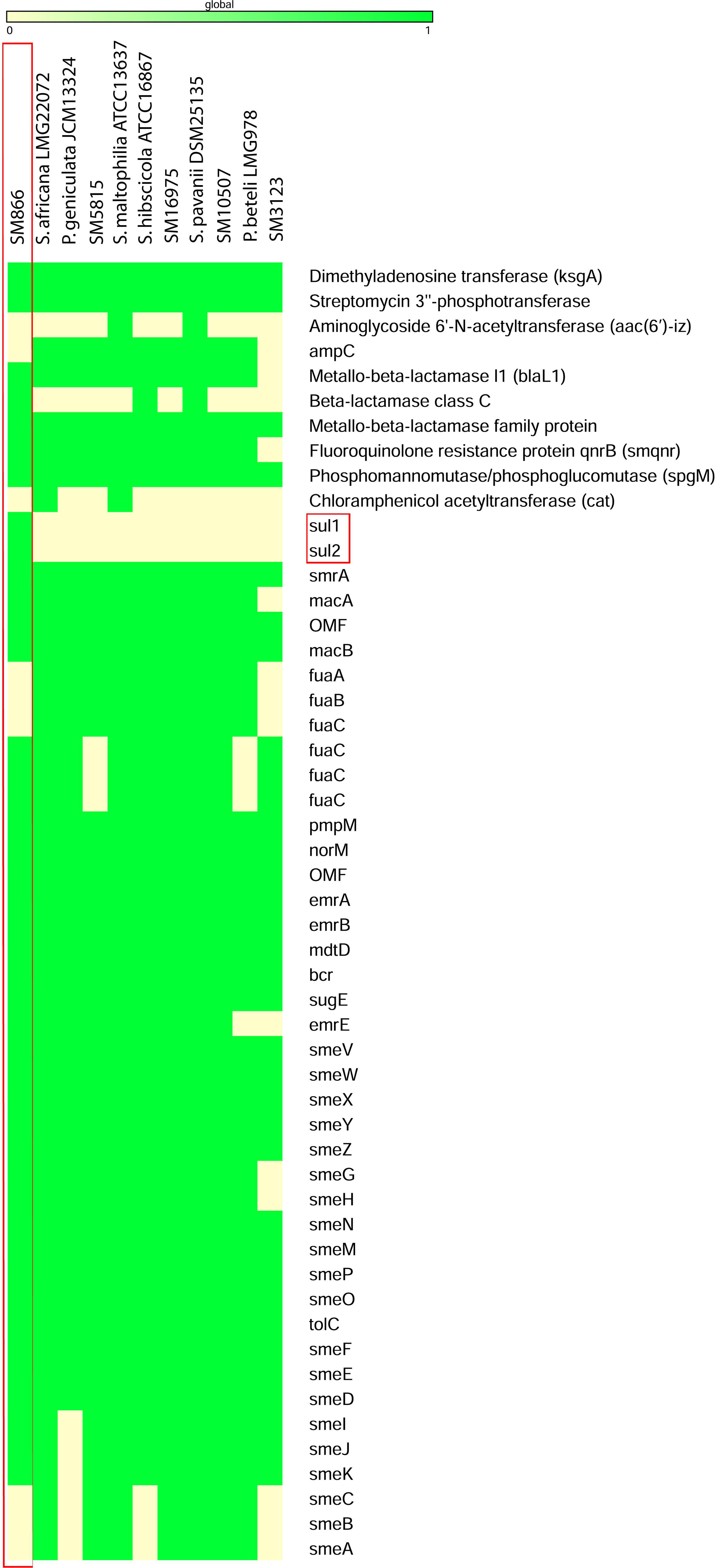
**Resistome heat map of Smc** distribution of antibiotic resistance and efflux pump genes across Smc. Genes and strains are indicated in x-axis and y-axis respectively. Here, presence and absence of gene is represented by green and yellow colors respectively. Here, SM866 is in highlighted in red box.

### Diverse super-integrons conferring extreme drug resistance

Genome mining of the SM866 revealed presence of two *sul* allelles (*sul1* and *sul2*) each on two distantly located integrons of chromosomal origin (figure 4(A)). To our knowledge, this is the first complete genome of *Stenotrophomonas* carrying *sul* genes. In addition to the *sul* genes, both the integrons were harbouring array of other drug resistance (tetracycline, phenicol, fluoroquinolone and macrolide antibiotics) and biocide resistance (*mer* operon, *qac* gene) genes linked to gene capturing machinery i.e. ISCR (IS91-like elements) element (figure 4(B)). Cassette I (3448861-3472206) (DUW70_16515 to DUW70_16640) was carrying *sul1*, truncated *qacE, bla(PME), aac(3), aph(3), floR2* and *tet(G)* genes in the ISCR. Cassette II (3052377-3082576) (DUW70_14405 to DUW70_14605) was carrying *sul2, floR, aph(3), aph(6)* genes in the ISCR along with *mer* operon. Additionally, SM866 is also having *smeDEF* (4366825-4372654) (DUW70_20855 to DUW70_20865) in the chromosomal region.

**Figure 4.**
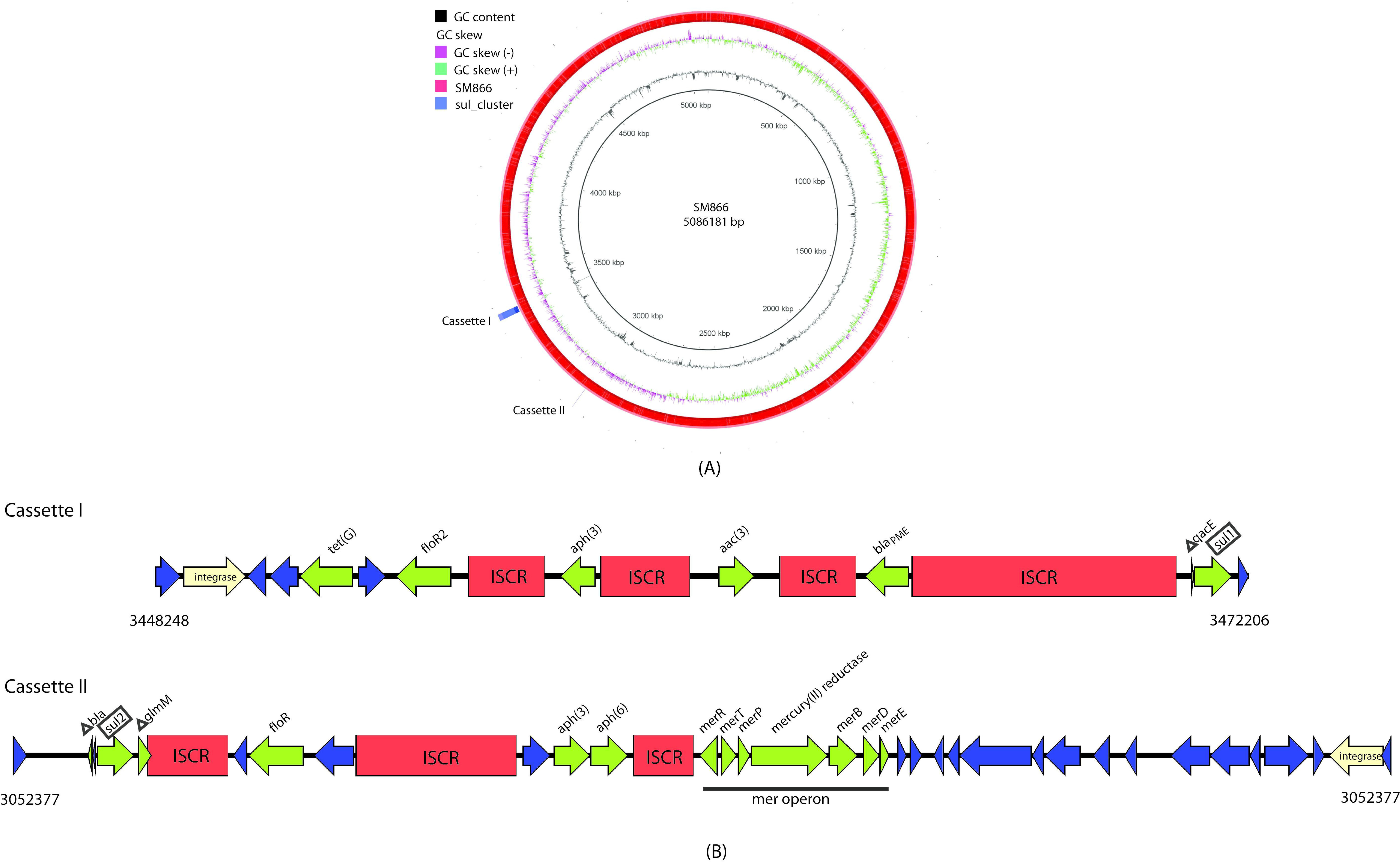
**(A) Circular representation of SM866** indicating chromosomal location of cassette I and cassette II. **(B) Schematic representation of antimicrobial genes in cassettes I and II** ISCR elements are shown in red color and antibiotic resistance genes are in green color.

## DISCUSSION

*S. maltophilia* complex strains demonstrate evolution of an opportunistic minor pathogen to novel genomospecies and can be a major threat in health care settings. Since *S. maltophilia* genomes displayed extreme fluidity at inter-strain level (35), there was need to understand extreme drug resistance of emerging strains at complete genome level. Till now, *S. maltophilia* has been studied at gene level (PCR-based) or draft genome level to understand extreme drug resistance (6–8). But, such studies are not giving the genomic context such as flanking region and origin of the resistance genes. In the present study, we have analyzed the complete genome of extreme drug resistant novel genomospecies of Smc. Our finding of two *sul* genes on two separate integrons in the chromosome indicates the systemic and rapid evolution of XDRs. This clearly depicts extreme selection pressure towards acquisition, maintenance and spread of multiple *sul* genes. At the same time, integrons carrying *sul* genes along with other commonly used antibiotics and biocide resistance in a highly evolved ISCR is alarming. Furthermore, till date, there is no complete genome available for a sulfamethoxazole resistant strain of Smc. This is also first complete genome sequence of an XDR strain from Smc complex (35). Hence, genome dynamics, considering the repetitive elements and mobilome of a strain resistant to main-stream drugs will be highly significant in understanding evolution of extreme drug resistance in the tertiary care units of Indian hospitals.

Genome dynamics is the key for survival and evolution of the new-age superbugs. Heavy usage of disinfectants like quaternary ammonium compounds and heavy metals is major activity in hospital settings (15). Yet, hospital acquired infections with multi-drug resistant bacteria are persistent and prevalent. Hence, the effectiveness of cleanliness is always under question as the hospital settings are shown to be rapidly contaminated by deadly pathogens with multidrug resistant organisms like: *Acinetobacter* spp., *Klebsiella pneumonia* etc. (36, 37). These nosocomial pathogens have been shown to persist in the hospital environment from some days to even months (38). In the present study, we have proposed that it is continuous biocide exposure, which might have favored the pathogen towards acquisition and maintenance of resistance through integron mediated rapid evolution. Since, ISCR element, along with antibiotic resistant genes, also carried genes for biocide and heavy metal resistance. Hence, the overdose of disinfectants and reduced susceptibility to them has acted as a potential selection for antibiotic resistance (39). Such co-evolution of antibiotic and biocide resistance has already been reported in literature for deadly pathogens (40, 41). Interestingly, biocide exposure of nosocomial pathogens have been reported to select for multidrug resistance (42).

Complete genome dynamics of SM866 revealed exclusive DRs encoding for various efflux pumps, mobile elements, transcriptional regulators etc. Multidrug efflux transporters of RND family, MFS family, MATE and ABC transporters families etc. have been already reported in *Salmonella enterica, Mycobacterium tuberculosis* contributing towards evolution of extreme drug resistance (43–45). In the present study, we are reporting similar condition for the first time in a novel genomospecies of *S. maltophilia* complex in context of chromosomally encoded exclusive dynamic regions. Furthermore, high patient density in developing country like India complicates the situation as seen by the potential of the integrons to rapidly diversify at chromosomal level and also to move across distant pathogens. Such a scenario results in evolution of complete drug resistance in nosocomial settings challenging their successful treatment.

Hence, in the present study, by investigation of the complete genome obtained using transformative long read technology, we are able to successfully reveal the cryptic evolutionary radiation in the emerging novel genomospecies of *S. maltophilia*. More such complete genome investigations of the superbugs at global level are required for a better understanding of the clinical risks of reduced susceptibility towards biocides, which has got little attention than antibiotic resistance. Further, present study emphasizes on awareness of targeted use of disinfectants restricting to the clinical benefit and not in indiscriminate manner.

## Conclusion

Complete genome sequencing allowed us to establish conclusively that *sul* genes are of chromosomal origin in our XDR strain. However, presence of two chromosomal integrons in a nosocomial strain indicated importance of chromosomal route and extent of selection leading to emergence of such a complex strain even in clinical settings. Further, Gillings noted in his review that chromosomal integrons in contrast to transient plasmid borne integrons, can become site to generate genomic and phenotypic complexity. In clinical setting, integrons are reported to be of plasmid origin and thought to be the result of human intervention (46). In this context, finding of a nosocomial pathogenic strain with two chromosomal integrons with complex genetic cassette suggests that clinical settings, like in environmental counterparts, have emerged as hotspots for integron evolution. Presence of antimicrobial genes in both integrons of a novel sequence type suggests nature of extreme selection happening in the hospital settings and further possibility of many more novel sequence types. Particularly, Indian hospitals are tropical clinical settings, having high patient density and heavy use of antimicrobials. Hence, a coordinated global research effort by integrating emerging long read technologies is needed for surveillance of usage of biocides for hygiene and linked/ rapid co-evolution of extreme drug resistance in nosocomial pathogens.

## Author statements

### Ethical approval

Ethics approval and each patient’s written consent was not required as it was a part of routine clinical testing.

## Abbreviations

XDR: extreme drug resistance
ISCR: IS91-like elements common regions
ST: sequence type
TMP/SXT: trimethoprim/sulfamethoxazole
DR: dynamic regions

## Supplementary data

**Supplementary figure 1(A) 16S rRNA phylogeny.** Here, SM 866 is in red box and Smc is in blue box. **(B) eBURST analysis of Smc.** Here, ST-366 is highlighted.

**Supplementary figure 2(A) Electrophoretic gel image** of plasmid screening with Fosmid as a positive control and *S. maltophilia* ATCC 13637 (Type strain) as negative control. Loading order: Lane1: λ/hind III ladder; Lane 2: Fosmid; Lane 3: Fosmid induced; Lane 4: *S. maltophilia* SM866; Lane 5: *S. maltophilia* ATCC13637; Lane 6: λ monocot ladder. **(B) Electrophoretic gel image** of the plasmid screen after treatment with ATP-dependent DNase. Loading order: Lane1: λ/hind III ladder; Lane 2: Fosmid; Lane 3: Fosmid (digested); Lane 4: Fosmid induced; Lane 5: Fosmid induced (digested); Lane 6: steno SM866; Lane 7: steno SM866 (digested); Lane 8: *S. maltophilia* ATCC13637; Lane 9: *S. maltophilia* ATCC13637 (digested); Lane 10: λ monocot ladder.

**Supplementary table 1: Dynamic regions** Genomic coordinates for the DRs exclusive to SM866.

