## Supplementary material for "Extreme genome selection towards complete antimicrobial resistance in a nosocomial strain of *Stenotrophomonas maltophilia* complex"

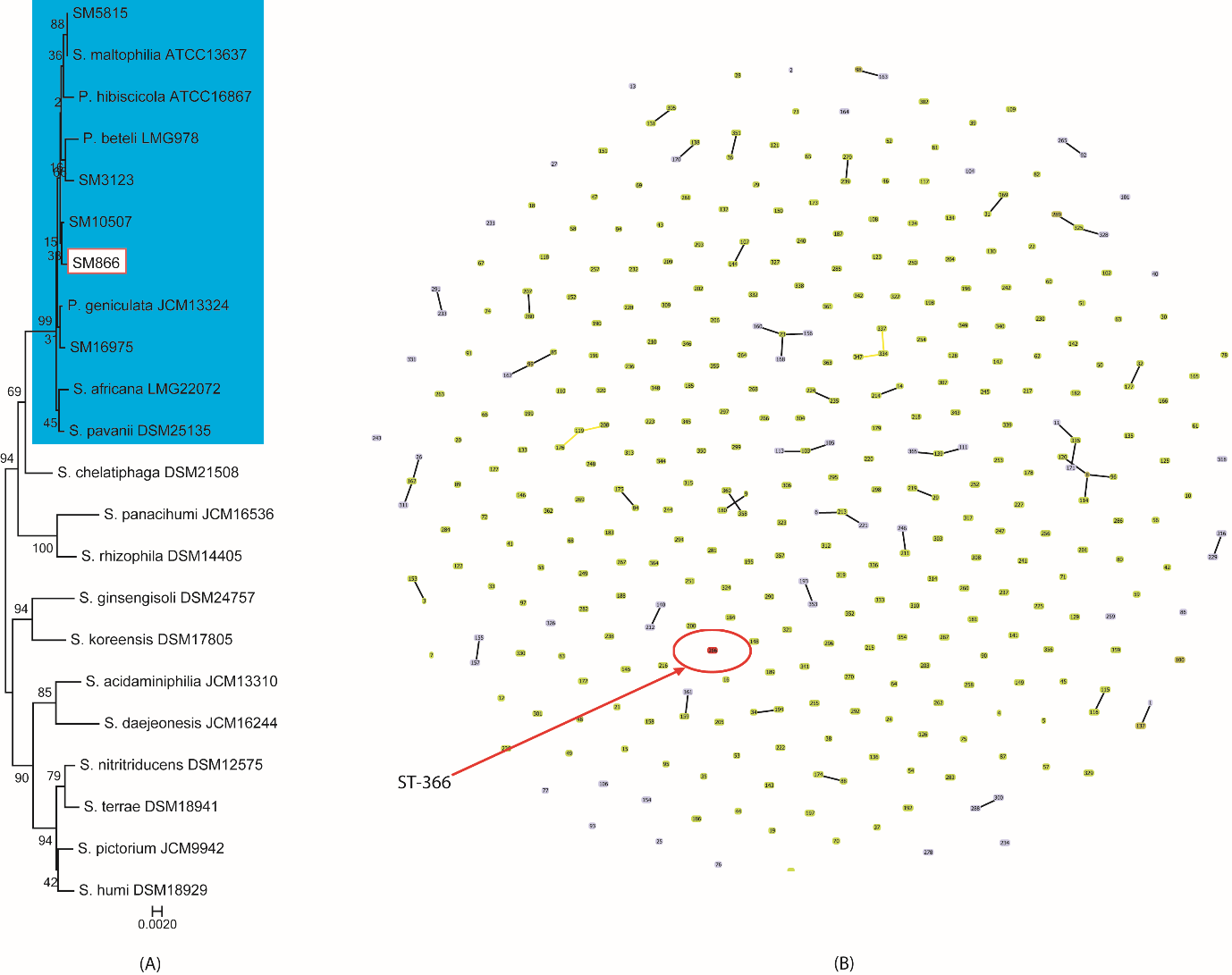

**Supplementary figure 1(A) 16S rRNA phylogeny.** Here, SM 866 is in red box and Smc is in blue box. **(B) eBURST analysis of Smc.** Here, ST-366 is highlighted.

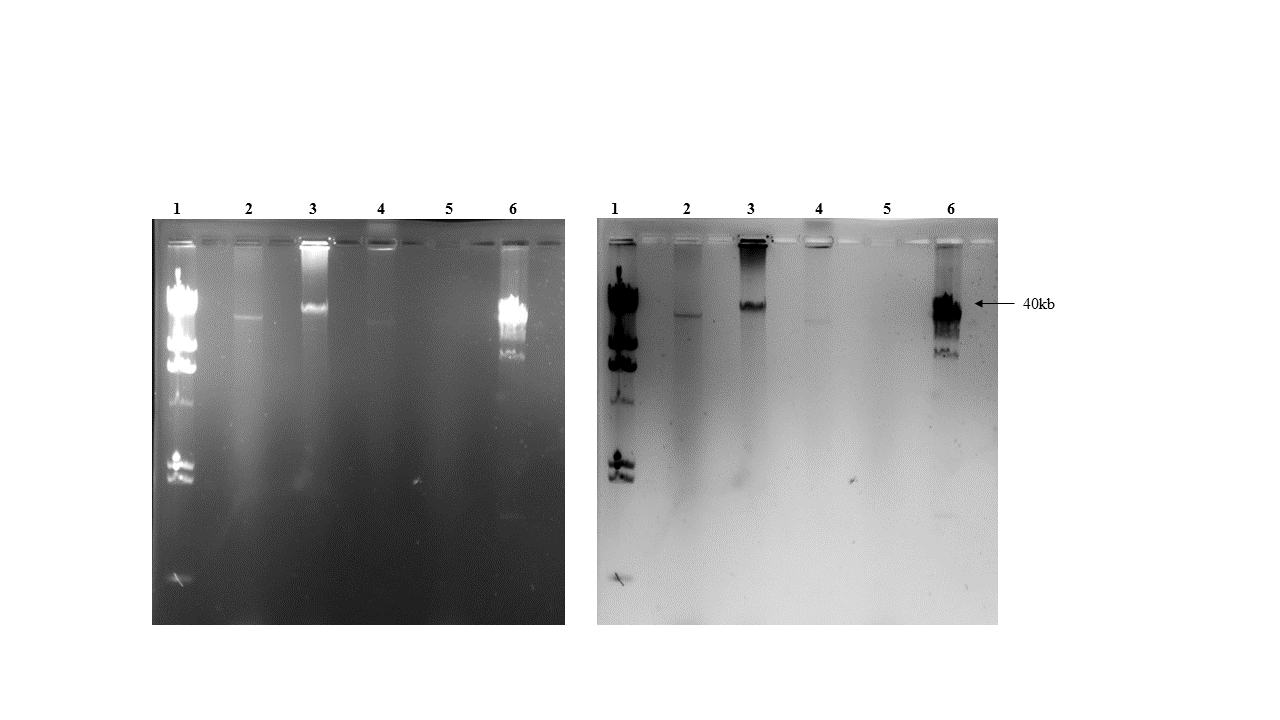

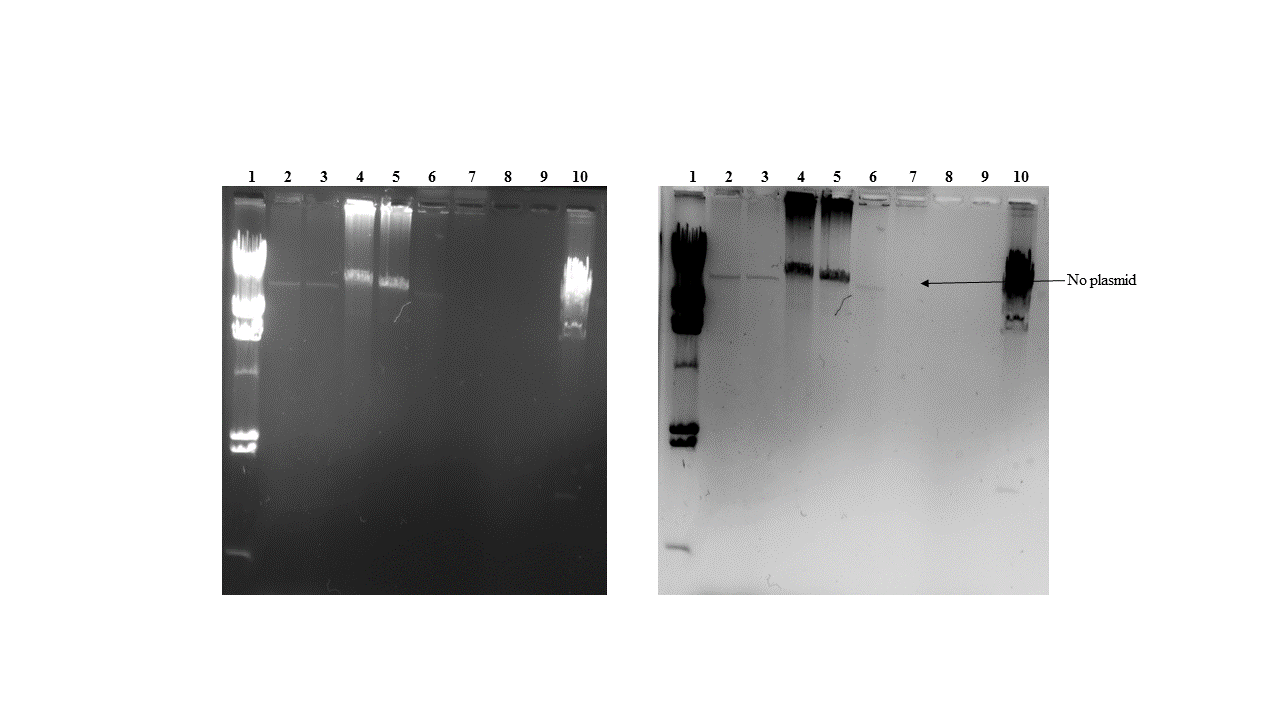

(A)

(B)

**Supplementary figure 2(A) Electrophoretic gel image** of plasmid screening with Fosmid as a positive control and *S. maltophilia* ATCC 13637 (Type strain) as negative control. Loading order: Lane1: λ/hind III ladder; Lane 2: Fosmid; Lane 3: Fosmid induced; Lane 4: *S. maltophilia* SM866; Lane 5: *S. maltophilia* ATCC13637; Lane 6: λ monocot ladder. **(B) Electrophoretic gel image** of the plasmid screen after treatment with ATP-dependent DNase. Loading order: Lane1: λ/hind III ladder; Lane 2: Fosmid; Lane 3: Fosmid (digested); Lane 4: Fosmid induced; Lane 5: Fosmid induced (digested); Lane 6: steno SM866; Lane 7: steno SM866 (digested); Lane 8: *S. maltophilia* ATCC13637; Lane 9: *S. maltophilia* ATCC13637 (digested); Lane 10: λ monocot ladder.

**Supplementary table 1: Dynamic regions** Genomic coordinates for the DRs exclusive to SM866.

| **Region** | **Start** | **End** | **Locus Tags** |
| --- | --- | --- | --- |
| **R1** | 159954 | 189692 | DUW70_00705 - DUW70_00840 |
| **R2** | 296193 | 326056 | DUW70_01330 - DUW70_01525 |
| **R3** | 401607 | 428898 | DUW70_01905 - DUW70_02085 |
| **R4** | 531295 | 540289 | DUW70_02540 - DUW70_02580 |
| **R5** | 657898 | 664003 | DUW70_03140 - DUW70_03155 |
| **R6** | 2048493 | 2084455 | DUW70_09745 - DUW70_09930 |
| **R7** | 2677242 | 2729455 | DUW70_12705 - DUW70_12970 |
| **R8** | 2981203 | 2996506 | DUW70_14080 - DUW70_14135 |
| **R9** | 3042076 | 3082345 | DUW70_14360 - DUW70_14605 |
| **R10** | 3184899 | 3245704 | DUW70_15065 - DUW70_15460 |
| **R11** | 3440153 | 3540552 | DUW70_16480 - DUW70_17075 |
| **R12** | 3639472 | 3672123 | DUW70_17540 - DUW70_17750 |
| **R13** | 4482792 | 4503612 | DUW70_21380 - DUW70_21445 |
| **R14** | 5064157 | 5071217 | DUW70_23945 - DUW70_23975 |
